# Human cytomegalovirus RNA2.7 regulates host cell cycle and facilitates viral DNA replication by inhibiting RNA polymerase II phosphorylation

**DOI:** 10.1101/378083

**Authors:** Yujing Huang, Jing Zhang, Xin Guo, Qing Wang, Zhongyang Liu, Yanping Ma, Ying Qi, Qiang Ruan

**Affiliations:** Virus Laboratory, Affiliated Shengjing Hospital, China Medical University, Shenyang, Liaoning, 110004, China; Department of Pediatrics, the Fourth Affiliated Hospital, China Medical University, Shenyang, Liaoning, 110032, China

**Author notes:** Corresponding author: Qiang Ruan.

## Abstract

Human cytomegalovirus (HCMV) is a ubiquitous pathogen belongs to the beta herpesvirus family. RNA2.7 is a viral long non-coding RNA accounting for more than 20% of total viral transcripts at early time of infection. By construction of RNA2.7 deleted mutant and genome transcriptomic analysis, RNA2.7 is demonstrated to repress host cellular RNA polymerase II (Pol II)-dependent transcription through inhibiting the phosphorylation of RNA polymerase II (Pol II). Co-immunoprecipitation, RNA immunoprecipitation and RNA electrophoretic mobility shift assay are followed to investigate its mechnism. A 145nt-in-length fragment in RNA2.7 is identified to bind to Pol II and block the interaction between Pol II and phosphorylated cyckin-dependent kinase 9 (phospho-CDK9). By inhibiting Pol II phosphorylation, RNA2.7 decreases the transcription and expression levels of chromatin licensing and DNA replication factor 1 (Cdt1) and cell division cycle gene 6 (Cdc6). Through above way, RNA2.7 prevents the entry of cells into S phase and facilitates viral DNA replication. Our results discover the functions of HCMV RNA2.7 in regulation of Pol II phosphorylation and cell cycle control during infection.

**Author summary:** Human cytomegalovirus (HCMV) RNA2.7 is a viral lncRNA that is most abundant during infection. Here we show that a 145nt-in-length fragment in RNA2.7 binds to RNA polymerase II (Pol II) and blocks the interaction between Pol II and phosphorylated cyckin-dependent kinase 9 (phospho-CDK9). By inhibiting Pol II phosphorylation, RNA2.7 decreases the transcription and expression levels of chromatin licensing and DNA replication factor 1 (Cdt1) and cell division cycle gene 6 (Cdc6), and blocks host cells entering into S phase. RNA2.7 is confirmed to facilitate viral DNA replication through decreasing Cdt1 and Cdc6. Therefore, our results discover the functions of HCMV RNA2.7 in regulation of Pol II phosphorylation and cell cycle control during infection.

## Introduction

Human cytomegalovirus (HCMV) is a ubiquitous human pathogen belongs to the beta herpesvirus family [1]. Following primary infection, the virus establishes lifelong latent infection with episodes of reactivation, mainly in the immune compromised host.

Long non-coding RNAs (lncRNAs) are roughly defined as RNA molecules of more than 200 bases in length without protein-coding capacity. More and more evidences indicate that lncRNAs may play an important role in a variety of biological processes and be involved in many human diseases including tumors and infections. Kaposi’s sarcoma-associated herpesvirus (KSHV) produces a highly abundant lncRNA known as PAN RNA. It has been demonstrated that PAN RNA controls KSHV gene expression by mediating the modification of chromatin through targeting KSHV repressed genome [2]. Recent high-resolution transcriptome mapping showed that most viral transcriptions during HCMV infection are concentrated in viral lncRNAs [3,4], including RNA1.2, RNA2.7, RNA4.9 and RNA5.0.

HCMV RNA2.7 is a viral lncRNA of 2.7-kb in length, which was previously termed beta2.7 [5,6]. RNA2.7 was observed to be abundant at early time of infection, accounting for more than 20% of total viral transcripts [7,8]. It has been found to prevent cell apoptosis by interacts with mitochondrial complex I during HCMV infection, and to be essential to maintain high levels of energy production in infected cells [9,10]. However, most functions of HCMV RNA2.7 involved in HCMV infection still remain unclear.

In this study, an activation of host cellular RNA polymerase II (Pol II)-dependent transcription was observed in cells infected with an RNA2.7 deleted HCMV strain. It was then identified that HCMV RNA2.7 could inhibit Pol II Serine2 (S2) phosphorylation by blocking phosphorylated cyckin-dependent kinase 9 (phospho-CDK9) binding to Pol II. The inhibition of Pol II S2 phosphorylation by HCMV RNA2.7 could decrease the transcription and expression levels of chromatin licensing and DNA replication factor 1 (Cdt1) and cell division cycle gene 6 (Cdc6) to regulate host cell cycle and facilitate viral DNA replication.

## Results

### An RNA2.7 deleted mutant is constructed based on HCMV bacterial artificial chromosome HAN

HCMV RNA2.7 is a transcript from the antisense strand in HCMV genome neighbored to RL1, RL6, RL8A and RL9A (Fig 1A). To study the functions of HCMV RNA2.7, an RNA2.7 deleted mutant (HANΔRNA2.7) was constructed based on a previously constructed HCMV bacterial artificial chromosome (BAC) HAN, which is the first characterized HCMV clinical strain in China [11]. By reverse-transcription PCR and sequencing, it was confirmed that the RNA2.7 sequence was deleted successfully in HANΔRNA2.7 genome (Fig 1B).

**Fig 1.**
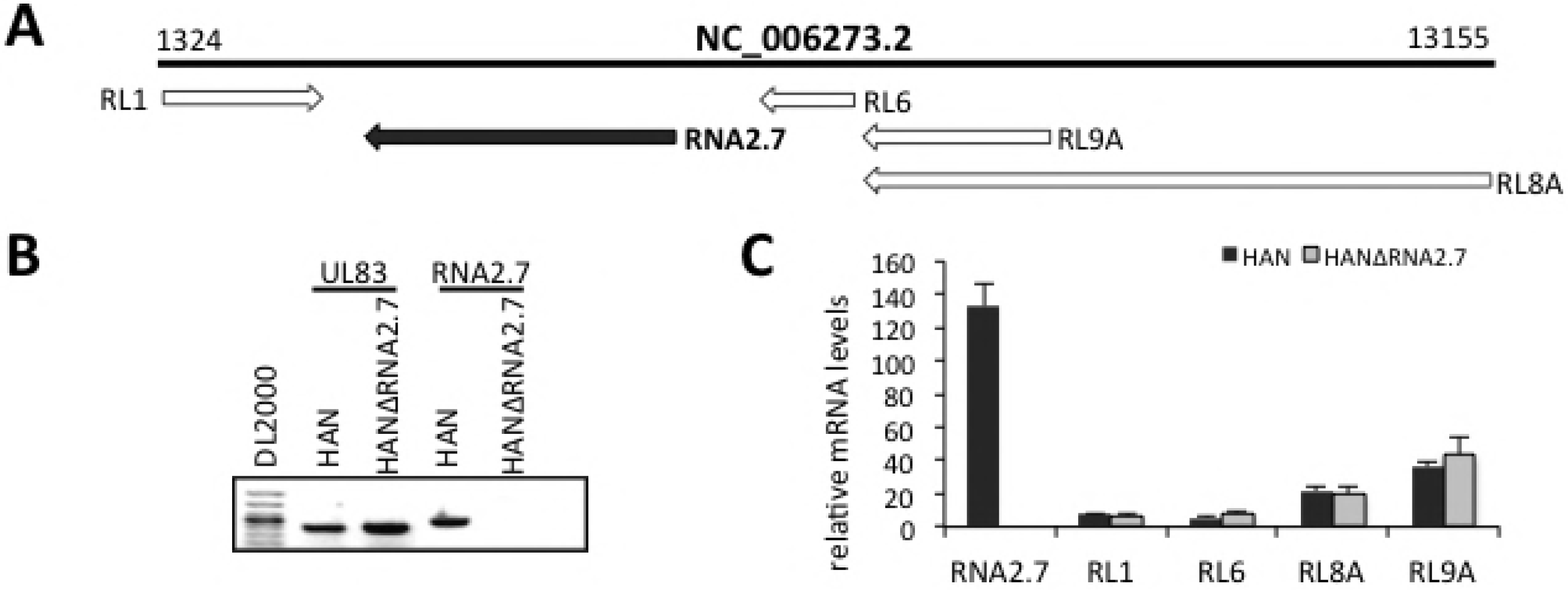
Construction of HCMV RNA2.7 deleted mutant. (A) Schematic diagram showing genomic location of HCMV RNA2.7. RNA2.7 is indicated in black bar. (B) Validation of RNA2.7 deletion. Reverse-transcription PCR for RNA2.7 transcript in HELF cells infected with HAN or HANΔRNA2.7. Transcript of HCMV UL83 was amplified as a control. (C) Quantitative PCR for RNA2.7 and its flanking genes in HELF cells infected with HAN or HANΔRNA2.7. Data are presented as mean±SEM.

To verify whether the deletion of RNA2.7 disturb the transcriptions of its flanking genes or not, human embryonic lung fibroblast (HELF) cells were infected with reconstituted viruses HAN and HANΔRNA2.7 at a multiplicity of infection (MOI) of 0.5, respectively. Total RNA was extracted at 72 hours post infection (hpi). The transcriptions of RNA2.7 and its flanking genes were measured by quantitative PCR respectively. Transcript of RNA2.7 was not detected in HANΔRNA2.7 infected cells (Fig 1C). The transcription levels of RL1, RL6, RL8A and RL9A in HANΔRNA2.7 infected cells were similar to those in HAN infected cells. Based on these results, HANΔRNA2.7 reconstituted virus could be used as an RNA2.7 knocked out HCMV stock in our further study.

### Host gene transcriptions are altered after infections with different HCMV constructs

To address the effects of RNA2.7 on host gene transcription, HELF cells were infected with HAN or HANΔRNA2.7 at an MOI of 0.5. HELF cells treated with PBS were used as a control. Total RNA was harvested at 72 hpi and subjected to whole genome transcriptomic analysis.

In total, transcriptions of 2,520 cellular genes were substantially altered by infection with HAN compared with uninfected cells, while transcriptions of 4,286 cellular genes were changed by HANΔRNA2.7 infection (Fig 2A). After sorting according to relative transcription levels, 4 patterns were identified: transcriptions of 509 genes were only affected by HAN infection and were designated as HAN specific transcriptional genes. Transcriptions of 54 cellular genes were inversely regulated between HAN and HANΔRNA2.7 infections, 1,957 genes were similarly regulated by both strains, and a total of 2,275 genes were HANΔRNA2.7 specific. In general, more cellular genes were transcribed in response to HANΔRNA2.7 infection compared with HAN infection, suggesting either an activation of gene transcription or a lack of suppression at transcriptional level due to deletion of RNA2.7.

**Fig 2.**
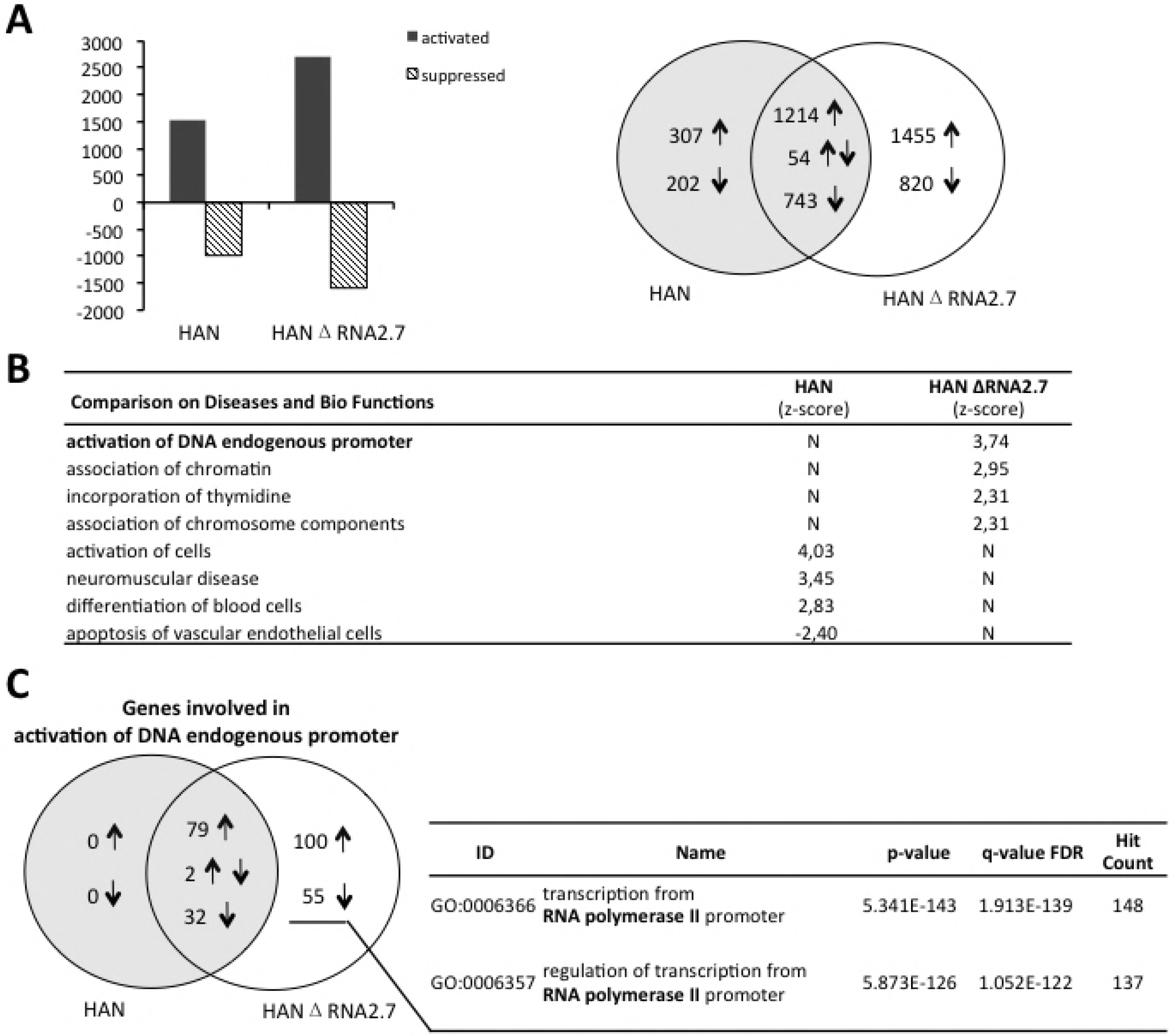
Repression of cellular Pol II-dependent transcription by RNA2.7. (A) General alteration of genome transcription in cells infected with HAN or HANΔRNA2.7. Cellular transcripts changed by infection are sorted in Venn diagram. (B) Comparison results on diseases and biofunctions of HAN or HANΔRNA2.7 altered transcripts. Their effects are indicated with z-score: an active effect is indicated with z-score>2.0; a suppressive effect is indicated with z-score<-2.0. (C) Venn diagram showing genes involved in activation of DNA endogenous promoter. HANΔRNA2.7 specific genes were analyzed and categorized into transcription from Pol II promoter.

### HCMV RNA2.7 represses cellular Pol II-dependent transcription

Cellular transcripts altered by HAN or HANΔRNA2.7 infection were analyzed for their comparisons on diseases and biofunctions. Results with different z-scores between HAN and HANΔRNA2.7 infected cells are listed (Fig 2B). In HAN infected cells, the apoptosis of endothelial cells was indicated to be repressed with z-score of -2.40. On the other hand, DNA endogenous promoter was indicated significantly activated in HANΔRNA2.7 infected cells (z-score=3.74). Genes involved in activation of DNA endogenous promoter were analyzed and their information is listed in S1 Table. No gene was grouped specific to HAN infection for DNA endogenous promoter activation. Except for 113 genes that their transcriptions were altered both in cells infected with HAN and HANΔRNA2.7, transcriptions of 155 genes associated with activation of DNA endogenous promoter were influenced specifically by HANΔRNA2.7 infection (Fig 2C). It was suggested that RNA2.7 might control host gene transcriptions through the suppression of DNA endogenous promoter activation, and the suppression was eliminated by deletion of RNA2.7.

The genes associated with activation of DNA endogenous promoter were further analyzed for gene ontology. It was predicted that 148 genes among the 155 genes specific to HANΔRNA2.7 infection (95.48%) were categorized into transcription from Pol II promoter. Based on these results, HCMV RNA2.7 was strongly indicated to play a role in the regulation of host Pol II-dependent transcription, suggesting an effect of RNA2.7 on quantity or activity regulation of Pol II during HCMV infection.

### HCMV RNA2.7 represses cellular Pol II-dependent transcription by inhibiting Pol II S2 phosphorylation

Pol II has been known to control most eukaryotic gene transcriptions including mRNA precursors and non-coding RNAs [12]. It is a multiprotein complex composed of 12 subunits. The C-terminal domain (CTD) of Pol II is a component of the largest subunit and contains multiple repeats of the heptapeptide consensus sequence YSPTSPS. Transcription through Pol II extensively depends on the phosphorylation of serine2 (S2) and serine5 (S5) in CTD.

To investigate the effect of HCMV RNA2.7 on Pol II, HELF cells were infected with HAN or HANΔRNA2.7. Total proteins were extracted at 0, 24, 48 and 72hpi respectively. Protein levels of Pol II and its two phosphorylated forms (Pol II S2 and Pol II S5) were measured by western blot. As shown in Fig 3A, Pol II S2 was significantly increased in cells infected with HANΔRNA2.7 compared to that in uninfected or HAN infected cells. The increase of Pol II S2 could be detected early at 24 hpi. However, Pol II and Pol II S5 proteins were not significantly different between HAN and HANΔRNA2.7 infected cells. Similar results were obtained by fluorescence microscopy detection. Pol II S2 in HANΔRNA2.7 infected cells was distributed in nucleus and higher than those in the HAN and mock infected cells (Fig 3B). It was demonstrated that the main effect of RNA2.7 on Pol II is to inhibit Pol II S2 phosphorylation during HCMV infection.

**Fig 3.**
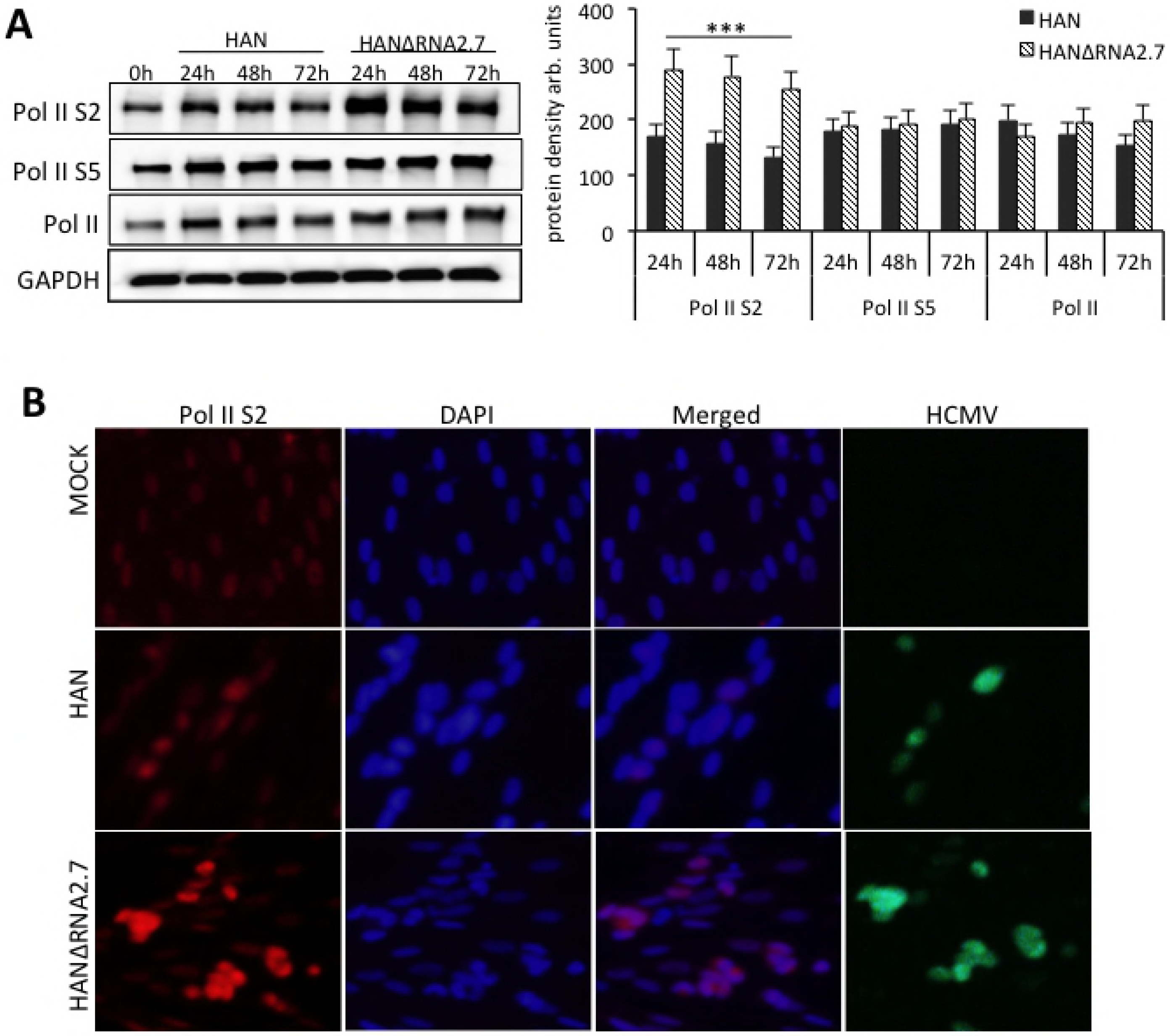
Inhibition of Pol II S2 phosphorylation by RNA2.7. (A) Western blot analysis for Pol II and its two phosphorylated forms (Pol II S2 and Pol II S5) in cells infected with HAN or HANΔRNA2.7 at different time points. The amounts of proteins are quantified by densitometry. Results are presented as mean±SEM. ***p<0.001. (B) Fluorescence microscopy detection of Pol II S2 in cells infected with HAN or HANΔRNA2.7. Pol II S2 protein was stained with rabbit anti-human Pol II S2 antibody and goat anti-rabbit IgG-TR. Nucleus was stained using ProLong Diamond Antifade Mountant with DAPI. Pol II S2 protein is shown in red and nucleus is shown in blue. HCMV virion with GFP is shown in green.

### HCMV RNA2.7 inhibits Pol II S2 phosphorylation by blocking interaction between phospho-CDK9 and Pol II

Phosphorylation of Pol II S2 promotes Pol II to overcome transcriptional blocks and increases the efficiency of 3’-end processing during elongation. Phospho-CDK9 is an important molecule mediating Pol II S2 phosphorylation. Therefore, protein levels of CDK9 and phospho-CDK9 were also measured and compared between cells infected with HAN or HANΔRNA2.7. Both CDK9 and phospho-CDK9 were increased and accumulated along with infection, while no difference was found between cells infected with HAN and HANΔRNA2.7 (Fig 4A). It was confirmed that inhibition of Pol II S2 phosphorylation by RNA2.7 was not directly caused by alteration of CDK9 or phospho-CDK9 proteins.

**Fig 4.**
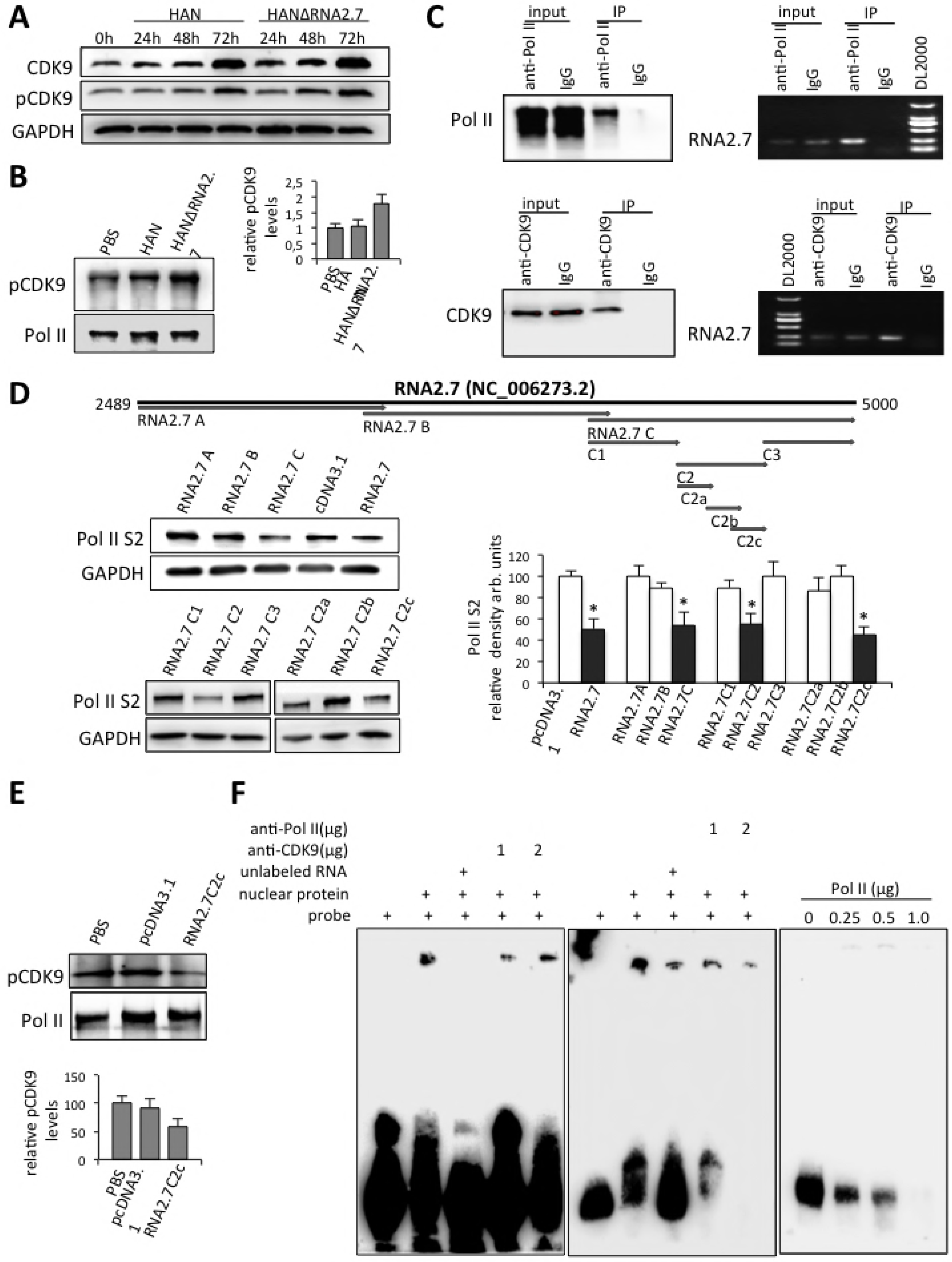
Block of interaction between Pol II and CDK9 by RNA2.7. (A) Western blot results for CDK9 and phosphorylated CDK9 (pCDK9) in cells infected with HAN or HANΔRNA2.7 at different time points. (B) Co-immunoprecipitation and western blot analysis of Pol II binding pCDK9 proteins in cells infected with HAN or HANΔRNA2.7. The relative amounts of pCDK9 binding to Pol II are quantified by densitometry with Pol II as a reference. Data are presented as mean±SEM. (C) Interaction between RNA2.7 and Pol II or CDK9. Pol II or CDK9 binding RNAs were immunopricipitated. Captured RNA was reverse transcribed and amplified using RNA2.7 specific primer. (D) Effect of RNA2.7C2c on inhibiting Pol II S2 phosphorylation. A series of vectors transcribing RNA2.7 fragments with different lengths were constructed and transfected into HELF cells. By western blot analysis, RNA2.7C2c was identified to inhibit Pol II S2 phosphorylation functionally. The amounts of Pol II S2 proteins are quantified by densitometry. Data are presented as mean±SEM. *p<0.05. (E) Effect of RNA2.7C2c on interaction between Pol II and pCDK9. Vector transcribing RNA2.7C2c was transfected into HELF cells. Cells transfected with pcDNA3.1 were used as a negative control. By co-immunoprecipitation and western blot analysis, RNA2.7C2c fragment was verified to block the interaction between pCDK9 and Pol II protein. The relative amounts of pCDK9 binding to Pol II are quantified by densitometry with Pol II as a reference. Data are presented as mean±SEM. (F) Confirmation of interaction between RNA2.7C2c fragment and Pol II protein by RNA EMSA.

Since the inhibition of Pol II S2 phosphorylation by RNA2.7 was not due to the quantity changes of CDK9 or phospho-CDK9, the effects of RNA2.7 on the interaction between phospho-CDK9 and Pol II was then studied. Pol II binding proteins were immunoprecipitated using anti-Pol II antibody in HELF cells infected with different strains. Phospho-CDK9 was measured in captured proteins by western blot. Pol II was used as a reference for calculation. When RNA2.7 was deleted during HCMV infection, phospho-CDK9 binding to Pol II was increased by more than 47.6% (Fig 4B). It was illustrated that RNA2.7 could block the binding of phospho-CDK9 to Pol II.

To explore the interaction between HCMV RNA2.7 and Pol II or CDK9, RNA immunoprecipitation (RIP) was performed to immunoprecipitate Pol II or CDK9 binding RNAs in HAN infected HELF cells. To exclude the disturbance of the binding between the undergoing transcriptional RNA2.7 and Pol II, alfa-amanitin, which can block all undergoing Pol II-dependent transcription processes, was added into cells 2 hours before immunoprecipitation. Captured RNA was reverse transcribed and amplified using RNA2.7 specific primers. RNA2.7 was found to bind to both Pol II and CDK9 proteins physically (Fig 4C).

### A 145nt fragment within RNA2.7 binds to Pol II and inhibits Pol II S2 phosphorylation

To investigate the functional RNA2.7 fragment mediating the inhibition of Pol II S2 phosphorylation, a series of vectors transcribing RNA2.7 fragments in different lengths were constructed and transfected into HELF cells (Fig 4D). Pol II S2 proteins in transfected cells were detected and compared at 48 hours post transfection. The Pol II S2 level in cells transfected with pcDNA3.1 was used as a reference. A fragment of 145nt (RNA2.7C2c) located in the 3’ terminus of RNA2.7 was identified to functionally inhibit Pol II S2 phosphorylation (Fig 4D). Compared to reference value, the level of endogenous Pol II S2 protein was decreased by 54.9% in cells transfected with vector transcribing RNA2.7C2c. The block effect of RNA2.7C2c was then validated using transfection and co-immunoprecipitation with anti-Pol II antibody. Compared to cells transfected with pcDNA3.1, the amount of phospho-CDK9 binding to Pol II was decreased by about 42.6% in cells transfected with vector transcribing RNA2.7C2c (Fig 4E).

To address the mechnism of RNA2.7C2c blocking the interaction between phospho-CDK9 and Pol II, RNA electrophoretic mobility shift assay (RNA EMSA) was carried out using biotin-labeled RNA2.7C2c RNA probes. A strip appeared after nucleoprotein was incubated with RNA2.7C2c probes (Fig 4F). The strip was weakened after competitive RNA or anti-Pol II antibody was added into the reaction systems. Although no classic super-shift strip was obtained after incubation with Pol II antibody, the strip was more weakened when more anti-Pol II antibody was added. However, no change of the strip was observed after incubation with anti-CDK9 antibody.

In addition, Pol II proteins were purified from nucleoproteins using magnetic beads and antibodies. Purified proteins with gradient increasing concentrations were incubated with biotin-labeled RNA2.7C2c RNA probes. When the concentration of purified Pol II protein was increased, the quantity of detected RNA probes was decreased correspondingly (Fig 4F). The results confirmed that RNA2.7C2c is a functional fragment of RNA2.7 to inhibit Pol II S2 phosphorylation by a physical binding to Pol II protein directly.

### HCMV RNA2.7 regulates host cell cycle by inhibiting Pol II S2 phosphorylation

Pathway analysis was carried out for cellular transcripts altered by HAN or HANΔRNA2.7 infection according to their known or suggested functions. Merged comparison was carried out then. After removal of RNA2.7 sequence from HCMV genome, pathways involved in cell cycle regulation were significantly influenced in infected cells (P<0.05), including cell cycle control of chromosomal replication and cell cycle regulation by BTG (B-cell translocation gene) family proteins (Fig 5A). A total of 11 genes related to cell cycle control of chromosomal replication, including Cdt1 and Cdc6, were increased in HANΔRNA2.7 infected cells, while none of them was changed in HAN infected cells.

**Fig 5.**
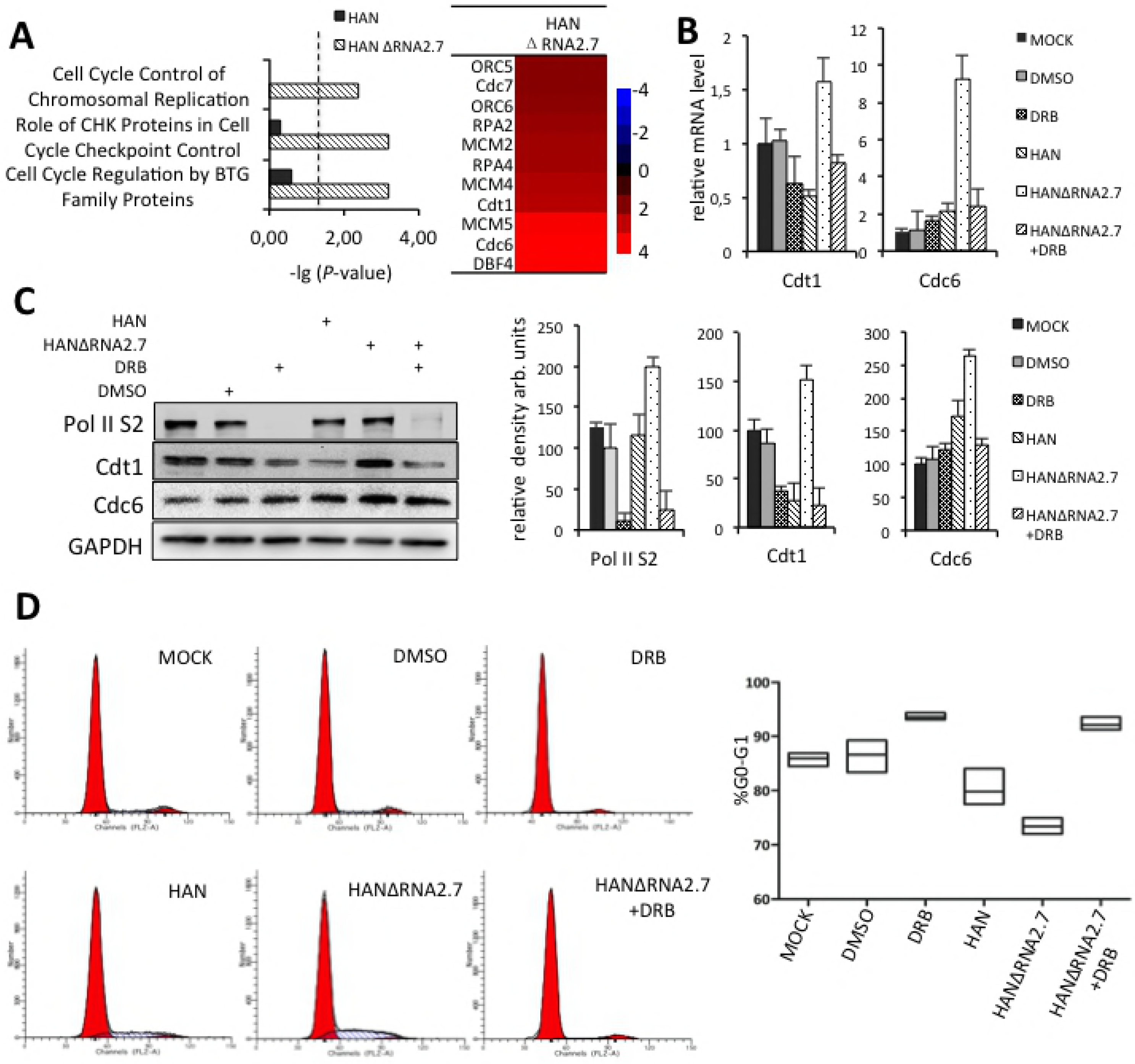
Regulation of host cell cycle by RNA2.7. (A) Results of pathway analysis indicating effects of RNA2.7 on pathways involved in cell cycle regulation. Heatmap shows that 11 genes involved in cell cycle control of chromosomal replication are increased in cells infected with HANΔRNA2.7. (B) and (C) Decreasing of mRNA and protein levels of Cdt1 and Cdc6 by RNA2.7 through inhibition of Pol II S2 phosphorylation. DRB was added 4 hours before infection with a final concentration of 200 nM and used as a positive control. Data are presented as mean±SEM. (D) Block of host cells entry into S phase by RNA2.7. Data are presented as mean±SEM.

To specifically address whether the changes in the cell cycle control of chromosomal replication was associated with the levels of Pol II S2 phosphorylation regulated by RNA2.7, 5,6-dichloro-1-b-D-ribofuranosyl benzimidazole (DRB) was used in this study. DRB is a specific inhibitor of Pol II S2 phosphorylation by blocking the interaction between phospho-CDK9 and Pol II proteins. HELF cells were pretreated using DRB at a final concentration of 200nM before HAN or HANΔRNA2.7 infection. The mRNA and protein levels of Cdt1 and Cdc6 were measured at 72 hpi. The results showed that both mRNAs and proteins of Cdt1 and Cdc6 were increased in HANΔRNA2.7 infected cells compared to those in HAN infected cells, and the effects were eliminated by DRB treatment (Figs 5B and 5C). It was speculated that HCMV RNA2.7 could decrease the transcription and expression of Cdt1 and Cdc6 by inhibiting Pol II S2 phosphorylation in the same manner with DRB.

Cdt1 and Cdc6 are important components of pre-replication complex. Formation of pre-replication complex could promote cellular DNA replication and drive the cells from G0/G1 phase into S phase. To investigate whether HCMV RNA2.7 could affect host cell cycle control through inhibiting Pol II S2 phosphorylation, the cell cycles were analyzed after different treatment and HCMV infection.

Compared with cells infected with HANΔRNA2.7, more cells (79.81% vs. 73.39%, *P*=0.0485) were arrested at G0/G1 phase when infected with HAN (Fig 5D). By inhibiting Pol II S2 phosphorylation with DRB, the cell population in G0/G1 phases of HANΔRNA2.7 infected cells was increased from 73.39% to 91.07% (*P*=0.0002). The results showed that HCMV RNA2.7 could decrease Cdt1 and Cdc6 levels and block host cells entering into S phase during HCMV infection through inhibiting Pol II S2 phosphorylation.

### HCMV RNA2.7 facilitates viral DNA replication by decreasing Cdt1 and Cdc6 expressions

Host cell cycle control is intimately linked to viral DNA replication. It was confirmed from our results that RNA2.7 could decrease expressions of Cdt1 and Cdc6 and regulate host cell cycle through inhibiting Pol II S2 phosphorylation. To study whether the repressing effects of RNA2.7 on Cdt1 and Cdc6 expressions could influence HCMV DNA replication during infection, Cdt1 and Cdc6 specific siRNAs were designed and transfected into HELF cells. Cdt1 and Cdc6 proteins were measured by western blot at 48 hours post transfection. The results showed that the expression of Cdt1 and Cdc6 could be effectively down regulated by specific siRNA transfection (Fig 6A).

**Fig 6.**
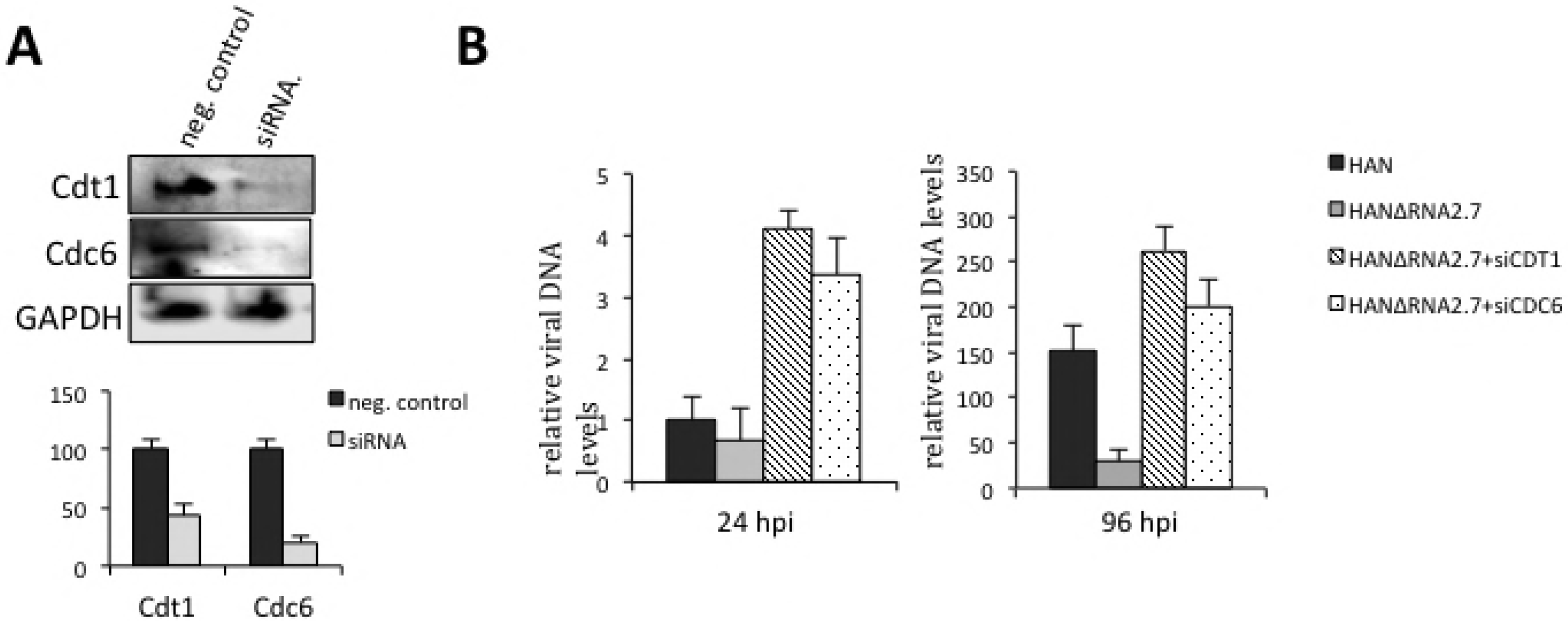
Effects of RNA2.7 on viral DNA replication. (A) Evaluation of siRNA Efficiencies specific to Cdt1 and Cdc6. The amounts of proteins are quantified by densitometry. Results are presented as mean±SEM. (B) Quantification of viral DNA at 24 and 96 hpi. Cells transfected with siRNAs specific to Cdt1 or Cdc6 were infected with HANΔRNA2.7. Cells transfected with siRNA negative control were infected with HAN or HANΔRNA2.7. Relative viral DNA levels are presented as mean±SEM.

HELF cells transfected with Cdt1 or Cdc6 specific siRNAs were infected with HAN or HANΔRNA2.7 at an MOI of 1.0. Relative viral DNA levels were measured at 24 and 96 hpi by quantitative PCR, respectively. The results showed that the removal of RNA2.7 sequence from viral genome caused a decrease of viral DNA replication by 33.63% at 24 hpi and 70.41% at 96 hpi. The supression of viral DNA replication by RNA2.7 deletion was reversed by inhibition of Cdt1 or Cdc6 expression using siRNAs. Compared to HANΔRNA2.7 infected cells, knock down of Cdt1 or Cdc6 during HANΔRNA2.7 infection resulted in a major increase with fold changes of 8.88 or 6.81 in viral DNA level at 96 hpi (Fig 6B). The results implied that HCMV RNA2.7 could facilitate viral DNA replication through decreasing Cdt1 and Cdc6 expressions.

## Discussion

Potential cellular protein partners that may interact with RNA2.7 directly were screened among cDNA library in lambda phage using RNA probe. NADH ubiquinone oxidoreductase, a subunit of mitochondrial complex I was identified to interact with RNA2.7. The interaction of RNA2.7 with mitochondrial complex I could inhibit rotenone-induced apoptosis in neuronal cells [9]. However, the functions of RNA2.7 are still far from clear.

To investigate the biofunctions of RNA2.7, genomic transcriptions were analyzed for cells infected with HCMV strain HAN or RNA2.7 deleted mutant HANΔRNA2.7 in our study. A general activation of cellular transcription was observed when RNA2.7 was removed. DNA endogenous promoter was indicated significantly activated in HANΔRNA2.7 infected cells and genes associated with activation of DNA endogenous promoter were categorized into transcription from Pol II promoter. The results indicate that RNA2.7 might repress cellular Pol II-dependent transcription during infection.

Viruses can extensively manipulate cellular gene expressions to maintain intracellular conditions benefiting viral survival and propagation. Transcription is the first step of gene expression and Pol II is an important enzyme participating in eukaryotic transcriptions. Diverse mechanisms of many viruses have been reported to regulate Pol II-dependent transcriptions. Herpes simplex virus I (HSV-1) has evolved redundant mechanism for triggering the loss of a Serine2-phosphorylated Pol II [13,14].

HCMV processes gene transcription through cellular Pol II and therefore it might also develop mechanisms to regulate Pol II activities contributing viral growth. It has been reported that HCMV UL79 interacts with Pol II to benefit accumulation of viral transcription during late stages of infection [15]. In our results, RNA2.7 inhibits the phosphorylation of Pol II S2 site without altering CDK9 and phospho-CDK9 levels. By a series of experiments including Co-immunoprecipitation and RNA immunoprecipitation, a 145nt-in-length fragment (RNA2.7C2c) in RNA2.7 was identified to inhibit Pol II S2 phosphorylation by blocking the interaction between phospho-CDK9 and Pol II.

The interaction between RNA2.7C2c and Pol II or CDK9 was detected using RNA EMSA. Different to adding anti-Pol II antibody into the reaction system, there was no change observed in the strips after adding anti-CDK9 antibody. No super-shift strip was obtained by adding anti-Pol II antibody into reaction system. It might be due to the large volume of RNA-protein complex that prevents them entering gel in electrophoresis. However, the detected quantity of biotin-labeled RNA2.7C2c reduced gradually along with the increasing input of purified Pol II correspondingly, which confirmed that the inhibition of Pol II S2 phosphorylation by RNA2.7C2c is mediated by a physical interaction between RNA2.7C2c and Pol II protein directly. The binding of RNA2.7C2c to Pol II might induce a competitive inhibition that reduces the binding of phospho-CDK9 to Pol II.

Deregulation of gene expression programs in cancer cells depends on continuous active transcription [16]. For example, the development of triple-negative breast cancer is mainly due to the uninterrupted transcription of oncogenes [17]. Disturbing transcription has been aimed for cancer therapy and chemical drugs triggers degradation of phosphorylated Pol II have been evaluated among cancer patients in recent years [18]. Transfection of small RNA molecule might cause less cytotoxicity than intake of chemical drugs. It has been reported that a small domain of RNA2.7 termed p137 can prevent dopaminergic cell death by protecting mitochondrial complex I activity and is expected to be used in therapy for Parkinson’s disease [10,19]. It might be similarly expected for RNA2.7C2c fragment to be used in clinical cancer therapy, considered to its effect of inhibiting Pol II S2 phosphorylation.

Cell cycle progression is regulated by a wide variety of factors. It is established that HCMV infection can lead to cell cycle arrest at some points [20,21]. The assembly of pre-replication complex at DNA replication origins during G1 phase of the cell cycle initiates cellular G1-S-phase transition. Cdt1 and Cdc6 are key components of cellular pre-replication complex [22–25]. In our study, both mRNA and protein levels of Cdt1 and Cdc6 were decreased along with the inhibition of POLR II S2 phosphorylation by RNA2.7. Without RNA2.7 during infection, more cells drove into S phase for cellular DNA replication.

Cellular DNA replication may compete against viral DNA replication, since more molecules are used for cellular DNA replication and less is available for viral DNA replication. Arresting host cells at G0/G1 phase is essential to the initiation of HCMV gene expression at the time of infection [26]. It has been reported that depletion of pre-replication factors in HCMV infected cell could promote viral replication [27]. Our data showed that HCMV DNA levels decreased when Cdt1 and Cdc6 were increased by removal of RNA2.7. The effect could be inverted by transfection with siRNA targeting Cdt1 and Cdc6.

Altogether, our results suggested a biological pathway by which RNA2.7 plays a role at the transcriptional level: HCMV RNA2.7 inhibits Pol II S2 phosphorylation by blocking the interaction between phospho-CDK9 and Pol II directly. The inhibition of Pol II S2 phosphorylation during infection decreases the transcriptions and expressions of Cdt1 and Cdc6, leads to host cell cycle arrest and facilitates viral DNA replication.

HCMV has both lytic and latent phases in its life cycle like other human herpesviruses. Although deletion of RNA2.7 did not result in growth defects of HCMV in fibroblast cells [28], the study on the functions of RNA2.7 should not be only limited in lytic phase of infection and permissive cells for lytic infection. RNA4.9 was proposed to play a role in transcriptional repression of viral IE gene expression for viral latency infection [4]. Similar to RNA4.9, RNA2.7 was detected in latent CD14+ and CD34+ cells at relatively high levels [4]. Based on our results, it is prospected that HCMV RNA2.7 might play a role in intracellular modulation in some aspects for the progress of latency or reactivation, such as cellular transcription and cell cycle control. More work on RNA2.7 functions for HCMV latency and reactivation is needed in future study.

## Materials and Methods

### Cells

Human embryonic lung fibroblast (HELF) cells were obtained from Shanghai Institutes for Biological Sciences (Cat#GNHu41). Cells were maintained in minimal essential medium (MEM) supplemented with 10% fetal bovine serum (FBS), 100 units/ml penicillin and streptomycin.

### Virus

Bacterial artificial chromosome of a characterized HCMV clinical low-passage isolate HAN (BAC HAN) has been constructed with green fluorescent protein (GFP) and identified previously (Zhao et al., 2016). BAC HAN stock was prepared on HELF cells and the virus titer was determined by standard TCID50 assays.

### Construction of HCMV RNA2.7 deleted mutant

RNA2.7 deleted mutant was constructed from BAC HAN. A fragment encoding kanamycin resistance gene was amplified from the previously described plasmid pGEM-oriV/Kan using forward and reverse RNA2.7Del primers (Forward: 5’-<u>AGATCGCTGCTGCTCCGGCGTTCTCCAGAAGCCCCGGCGGGCGAATCGGC</u>CTGTCTCTTATACACATCTCAACCATC-3’; Reverse: 5’-<u>GCATGCAAACTTCTCATTTATTGTGTCTACTACTCTGTGTTGCTACAGGGAG</u>CTGTCTCTTATACACATCTCAACCCTG-3’; Sequence homologous to RNA2.7 flanking regions was underlined). Purified PCR product was electro-transferred into competent bacteria DY380 that contains BAC HAN construct. Colonies were picked and the deletion of HCMV RNA2.7 was approved by PCR and sequencing directly.

BAC HANΔRNA2.7 plasmid was extracted from the identified bacteria DY380 colony using NucleoBondXtra Midi Kit (Macherey-Nagel) following the manufacturer’s protocol and transfected into HELF cells by electroporation. The cells were then maintained under standard cell culture conditions until cytopathic effects (CPEs) and GFP signal were clearly apparent. The reconstituted virus was named as HANΔRNA2.7, and the virus stock was prepared as described previously.

### Validation of HCMV RNA2.7 deletion

HELF cells growing in 6-well plates were infected with HAN or HANΔRNA2.7 at an MOI of 0.5 for 72 hours. Total RNAs were extracted from cells using Trizol reagent (ThermoFisher) according to the protocol, dissolved in 40μl RNase free water and then treated by TURBO DNA-free™ Kit (ThermoFisher). RNA preparations were estimated by electrophoresis on 1% agarose gel and quantified by ND-1000 spectrophotometer (Nanodrop Technologies). Total RNAs were reverse-transcribed using the SuperScript III First-strand synthesis system (ThermoFisher).

Primers RNA2.7rt forward and reverse were designed and used to amplify RNA2.7 from the cDNAs. Transcript of HCMV UL83 was amplified as a control. The primer sequences used in reverse-transcription PCR are shown in Table 1. Products were analyzed by electrophoresis on a 1.5% agarose gel containing ethidium bromide and visualized under ultraviolet light.

**Table 1.**
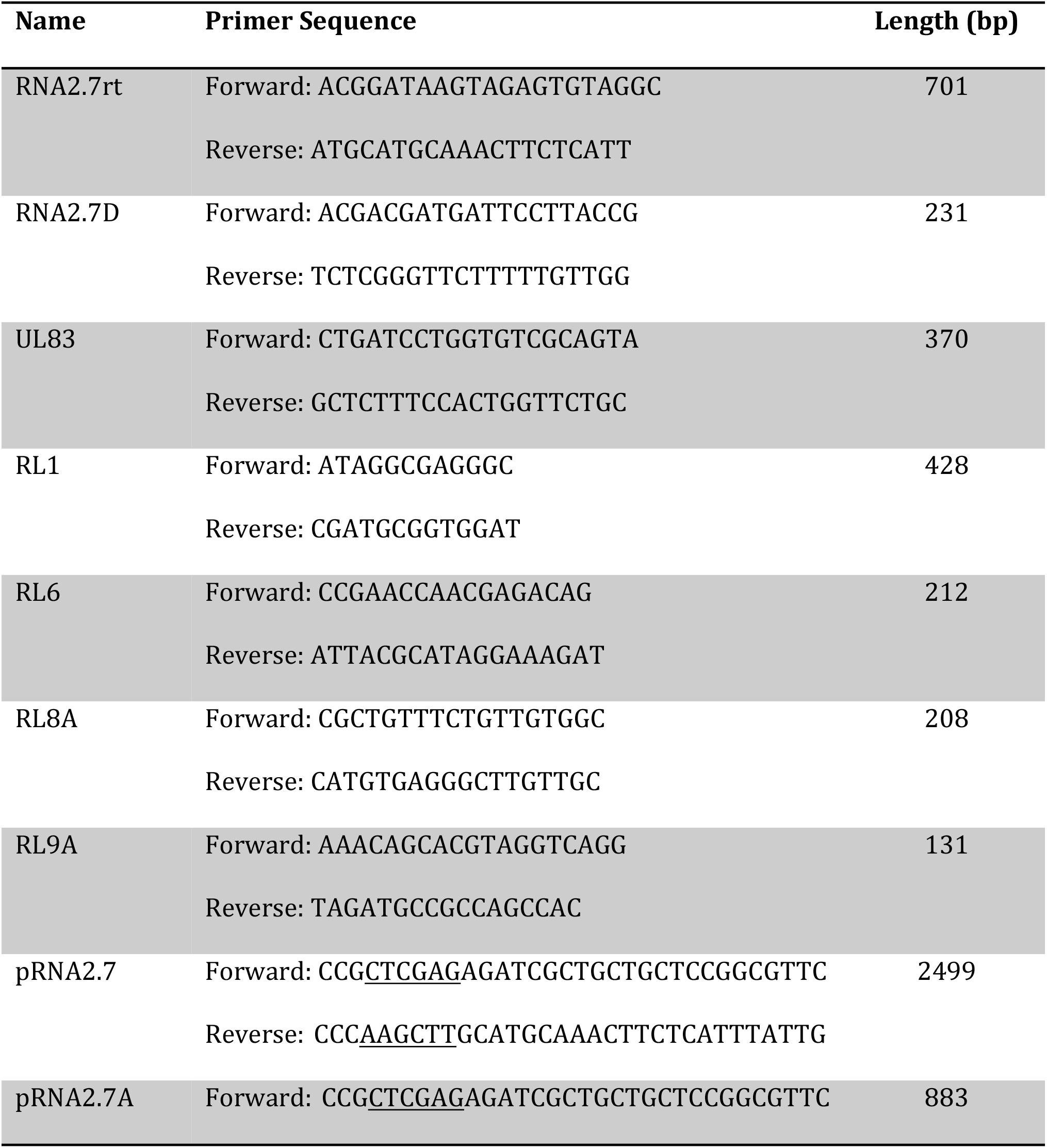
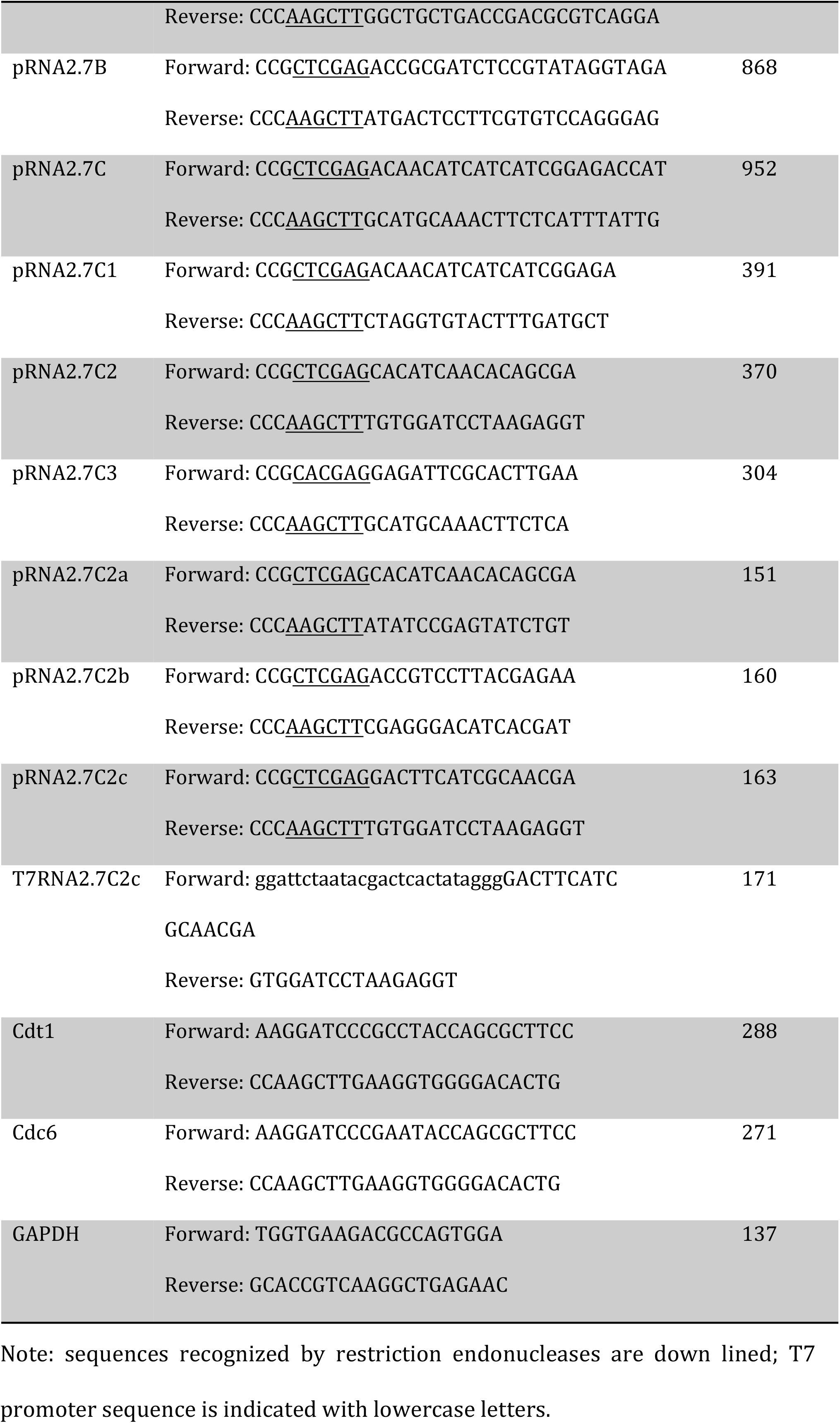
Sequences of primers used in this study

The transcripts of HCMV RNA2.7 and its flanking genes were quantified using QuantiTect SYBR Green-based PCR Kits (Qiagen) on Applied Biosystems 7300 Fast Real-Time PCR System (Applied Biosystems). Primers for quantitative PCR are listed in table 1. Detections were performed in biological triplicates and the relative transcription levels of RNA2.7, RL1, RL6, RL8A and RL9A were normalized to that of housekeeping gene glyceraldehyde 3-phosphate dehydrogenase (GAPDH) in corresponding samples using 2^-ΔΔCT^ methods.

### Transcriptomic microarray assays and statistical analysis

HELF cells growing in 100mm plates were infected with HAN or HANΔRNA2.7 at an MOI of 0.5. Total RNAs was extracted and purified at 72 hpi using RNeasy Mini Kit (Qiagen) following the manufacturer’s instruction and checked to inspect RNA integration by an Agilent Bioanalyzer 2100 (Agilent technologies). Microarray assays were performed at Shanghai Bio Corporation (National Engineering Center for Biochip in Shanghai, China) using GeneChipTM Human Genome U133 Plus 2.0 Assays with 54,675 probes representing 29,255 genes/transcripts (ThermoFisher). Raw data were processed and normalized by Expression Console software (version 1.1.2). Genes with empirical Bayes-adjusted *P* values less than 0.05 and fold change greater than 2.0 were considered differentially transcribed.

The microarray data are available in the NCBI’s Gene Expression Omnibus database (accession number GSE73954). Differentially transcribed genes were functionally categorized using ToppGene (https://toppgene.cchmc.org) and Ingenuity Pathway Analysis software (Ingenuity Systems).

### Western blot analyses of Pol II and CDK9

HELF cells growing in 6-well plates were infected with HAN or HANΔRNA2.7 at an MOI of 0.5. Cells were harvested and re-suspended using mammalian protein extraction reagent M-PER (ThermoFisher) with protease and phosphatase inhibitor cocktail (Abcam). Lysates were separated by SDS-PAGE. Blotted PVDF membranes were incubated with antibodies against Pol II, Pol II S2, Pol II S5, CDK9, phospho-CDK9 and GAPDH, followed by anti-IgG-HRP with ECL western blot reagent (ThermoFisher). Protein relative densities were quantified using ImageJ software 1.44p (NIH). GAPDH was used as a quantitative control.

### Pol II S2 staining and microscopy detection

HELF cells were cultured on NuncLab-Tek 8 well chamber glass slides (ThermoFisher) and infected with HAN or HANΔRNA2.7. Cells treated with PBS were used as a control. At 72 hpi, cells were washed, permeabilized and then stained with rabbit anti-human Pol II S2 antibody (Abcam) and goat anti-rabbit IgG-TR (Santa). Slides were washed and left to dry. Nucleus was stained using ProLong Diamond Antifade Mountant with DAPI (ThermoFisher). Stained cells were viewed and counted under fluorescence microscopy. Images were analyzed using Imag J software 1.44p (NIH).

### Vector construction and transfection

HCMV RNA2.7 sequence was amplified from cDNA using PrimerSTAR Max DNA Polymerase Mix (TaKaRa). A series of RNA2.7 fragments in different lengths were obtained using different primers. The primers used in the vector construction are listed in Table 1. PCR products were inserted into pcDNA3.1 (-) vector (ThermoFisher) and transformed into TOP10 (TianGen BioTech). All constructs were verified by sequencing.

HELF cells were prepared in 6-well plates. 2μg vectors per well were transfected into the cells using Attractene Transfection reagent (Qiagen) after cell preparation. Cells without transfection or transfected with pcDNA3.1 (-) vector were used as a negative control. 48 hours after transfection, proteins were extracted for western blot analysis or Co-IP.

### Co-immunoprecipitation

HELF cells growing in 100mm plates were infected with different HCMV strains at an MOI of 0.5 or transfected with vectors transcribing RNA2.7C2c. Cells were harvested and re-suspended using M-PER (ThermoFisher) with protease and phosphatase inhibitor cocktail (Abcam). PureProteome™ProteinA/G Mix Magnetic Beads (Millipore) were coated with anti-Pol II antibody and incubated with lysates at 4°C overnight. The captured protein complex was eluted with 60μl SDS-PAGE sample loading buffer (Beyotime) and then heated at 70°C for 10 minutes. After the beads were removed, the supernatants were loaded on 8% SDS-PAGE gel. Blotted PVDF membranes were incubated with antibodies against Pol II and phospho-CDK9, followed by peroxidase-conjugated goat anti-mouse or rabbit IgG (ZSGB-BIO) with ECL western blot reagent (ThermoFisher). Protein relative densities were quantified using ImageJ software 1.44p (NIH). Pol II was used as a quantitative reference to caculate the relative amount of phospho-CDK9 binding to Pol II.

### RNA immunoprecipitation

1×10^7^ HELF cells were infected with HAN at an MOI of 0.5 for 72 hours. Cells were treated with alfa-amanitin (MedChem Express) 2 hours before cell collection at a final concentration of 1 μg/ml. RIP experiments were performed using the Magna RIP RNA-Binding Protein Immunoprecipitation Kit (Millipore). Cells were pelleted and re-suspended with RIP lysis buffer plus protease and RNase inhibitors. The cell lysates were incubated with rabbit anti-CDK9 antibody (Abcam) or rabbit anti-Pol II antibody (Millipore) coated beads overnight according to the protocol. Rabbit IgG was used as a control. After treating with proteinase K, the immunoprecipitated RNAs were extracted by RNeasyMinElutte Cleanup Kit (Qiagen) and reversely transcribed using PrimeScript RT Master Mix (TaKaRa). The abundance of RNA2.7 was detected using primers RNA2.7D forwards and reverse by RT-PCR. Primer information is listed in table 1.

### Preparations of nucleoprotein and Pol II protein

Nucleoprotein was extracted from 1×10^7^ HELF cells using NE-PER Nuclear and Cytoplasmic Extraction Reagents (ThermoFisher) according to its instrument. Pol II protein was purified using PureProteome™Protein A/G Mix Magnetic Beads (Millipore) and anti-Pol II antibody (Millipore). The captured Pol II protein was eluted with 60μl native elution buffer (0.2M Glycine-HCl, PH 2.5) and then neutralized by adding 5μl neutralization buffer (1M Tris-HCl, PH 8.5).

### RNA electrophoretic mobility shift assay

RNA2.7C2c sequence containing T7 promoter was generated by PCR using Phusion High-Fidelity PCR Master Mix (NEB). PCR products were cloned and sequenced. RNA2.7C2c probe and competitive RNA2.7C2c were transcribed using T7 RNA Polymerase (NEB). RNA2.7C2c probe was labeled with biotin by adding biotin-16-UTP (Roche) into transcription reaction. After DNase treatment, probes were purified using RNeasy Mini Kit (Qiagen) and kept at -80°C till use.

RNA EMSA was carried out with LightShift^®^Chemiluminescent RNA EMSA Kit (ThermoFisher). To relax RNA folding, probes were heated at 80°C for 5 minutes and then placed on ice before use. Binding reactions were assembled with 2nM RNA2.7C2c probe and 4μg nucleoprotein or purified Pol II protein with gradient concentrations in binding buffer, 5% glycerol and 25nM DTT in 20μl. In addition, tRNA (0.5μg/μl final concentration) was used as a nonspecific competitor and unlabeled RNA2.7C2c was used as specific competitor. Anti-Pol II antibody (Abcam) was added for super shift. After 20 minutes incubation at room temperature, the samples were loaded onto 4% polyacrylamide gel in TBE buffer and run for 90 minutes at 100 volts at 4°C. Samples were transfered to Nylon membrane at 400mA for 60 minutes and crosslinked using a commercial ultraviolet light crosslinking instrument. Biotin-labeled probes were detected with Chemilumiescent Nucleic Acid Detection Module (ThermoFisher).

### Quantitative PCR and western blot analyses of Cdt1 and Cdc6

HELF cells growing in 6-well plates were infected with HAN or HANΔRNA2.7 at a MOI of 0.5 for 72 hours. DRB was added into the supernatant 4 hours before cell collection at a final concentration of 200nM. Mock infected cells and cells treated with DMSO were used as controls.

Total RNA samples were extracted and reverse-transcribed using the SuperScript III First-strand synthesis system (ThermoFisher). Transcriptions of Cdt1 and Cdc6 were quantified from the cDNA using QuantiTect SYBR Green-based PCR Kits (Qiagen) on Applied Biosystems 7300 Fast Real-Time PCR System (Applied Biosystems). Primer details are listed in table 1. The reaction conditions were as follows: an initial denaturation at 95°C for 2 minutes, then 40 cycles of annealing/extension at 60°C for 30 seconds followed by a final denaturation at 95°C for 15 seconds. Detections were performed in biological triplicates and the relative transcription levels of Cdt1 and Cdc6 were normalized to that of housekeeping gene GAPDH in corresponding samples using 2^-ΔΔCT^ method.

Protein samples were extracted using mammalian protein extraction reagent (ThermoFisher) with protease and phosphatase inhibitors. Lysates were separated by SDS-PAGE. Blotted PVDF membranes were incubated with antibodies against Pol II S2, Cdt1, Cdc6 and GAPDH, followed by peroxidase-conjugated goat anti-mouse or rabbit IgG (ZSGB-BIO) with ECL western blot reagent (ThermoFisher). Protein relative densities were quantified using ImageJ software 1.44p (NIH) using GAPDH as a control.

### Propidium iodide staining

HELF cells growing in 25T-flasks were infected with HAN or HANΔRNA2.7 at an MOI of 0.5 for 72 hours. Four hours before harvest, the HANΔRNA2.7 infected or uninfected cells were treated with DRB at a final concentration of 200nM. Uninfected cells and cells treated with DMSO were used as controls. Cells were washed twice in ice cold PBS then harvested using cell dissociation buffer before being centrifuged. Cell pellets were resuspended in 70% ethanol and kept at -80°C overnight. The fixed cells were then washed in PBS and resuspended in PI/RNase Staining buffer (BD). Cells were incubated for 30 minutes at room temperature, washed twice in PBS, resuspended in 0.5ml PBS and finally analyzed by flow cytometry. Single cells were gated and percentages of cells in the G0/G1 phases of the cell cycle were calculated using winMDi.

### SiRNA transfection

siRNAs specific to Cdt1 and Cdc6 (RiboBio) were diluted with RNase free water to a final concentration of 20μM. All siRNAs were stored as aliquots at -80°C to avoid multiple freeze thaw cycles. The siRNA sequences are as follows: siRNA to CDT1: GCACCAGGAGGT CAGATTA; siRNA to Cdc6: GTGTGAGACTATTCAAGCA.

HELF cells were prepared in 6-well plates with 70% confluence. Transfection of siRNAs was carried out using HiPerfect Transfection Reagent (Qiagen) according to its instrument. The final concentration of siRNA in each well was 100nM. MEM with 10% FBS was changed by MEM with 2% FBS at 24 hours post transfection for viral infection.

### Quantitative PCR of viral DNA

SiRNAs transfected HELF cells were infected with HAN or HANΔRNA2.7 at an MOI of 0.5. DNA samples were extracted at 24 and 96 hours post infection respectively. Quantitative PCR was carried out for HCMV UL83 and GAPDH using QuantiTect SYBR Green-based PCR Kits (Qiagen) on Applied Biosystems 7300 Fast Real-Time PCR System (Applied Biosystems). Primer details are listed in table 1. The reaction conditions were as follows: an initial denaturation at 95°C for 2 minutes, then 40 cycles of annealing/extension at 60°C for 30 seconds then denaturation at 95°C for 15 seconds. Detections were performed in biological triplicates and the relative level of viral DNA was normalized to that of housekeeping gene GAPDH in corresponding samples using 2^-ΔΔCT^ methods.

### Statistical analysis

Statistical analyses were performed using Excel and GraphPad Prism 5.0. Statistically significant differences were calculated using unpaired 2-tailed Student’s *t*. In all cases, *P* value less than 0.05 were considered significant.

## Acknowledgements

We thank all the colleagues mentioned in Materials and Methods for providing important reagents.

## Supporting Information

**S1 Table.** Genes involved inactivation of DNA endogenous promoter

## References

1. Landolfo S., Gariglio M., Gribaudo G., Lembo D. The human cytomegalovirus. Pharmacol Ther. 2003; 98:269–297.

2. Rossetto C.C., Pari G.S. Kaposi’s sarcoma-associated herpesvirus noncoding polyadenylated nuclear RNA interacts with virus- and host cell-encoded proteins and suppresses expression of genes involved in immune modulation. J. Virol. 2011; 85:13290–13297. doi: 10. 1128/JVI.05886-11.

3. Gatherer D., Seirafian S., Cunningham C., Holton M., Dargan D.J., Baluchova K, et al. High-resolution human cytomegalovirus transcriptome. Proc Natl Acad Sci USA. 2011; 108:19755–19760.

4. Rossetto C.C., Tarrant-Elorza M., Pari G.S. Cis and trans acting factors involved in human cytomegalovirus experimental and natural latent infection of CD14 (+) monocytes and CD34 (+) cells. PLOS Pathog. 2013; 9(5): e1003366.

5. McDonough S.H., Spector D.H. Transcription in human fibroblasts permissively infected by human cytomegalovirus strain AD169. Virology. 1983; 125: 31–46.

6. McDonough S.H., Straprans S.I., Spector D.H. Analysis of the major transcripts encoded by the long repeat of human cytomegalovirus strain AD169. J Virol. 1985; 53: 711–718.

7. Greenaway P.J., Wilkinson G.W. Nucleotide sequence of the most abundantly transcribed early gene of human cytomegalovirus strain AD169. Virus Res. 1987; 7(1): 17–31.

8. Spector D.H. Activation and regulation of human cytomegalovirus early genes. Intervirology. 1996; 39 (5-6): 361–377.

9. Reeves M.B., Davies A.A., McSharry B.P., Wilkinson G.W., Sinclair J.H. Complex I binding by a virally encoded RNA regulates mitochondria-induced cell death. Science. 2007; 316 (5829): 1345–1348.

10. Kuan W.L., Poole E., Fletcher M., Karniely S., Tyers P., Wills M., et al. A novel neuroprotective therapy for Parkinson’s diaease using a viral noncoding RNA that protects mitochondrial complex I activity. J Exp Med. 2012; 209 (1): 1–10.

11. Zhao F, Shen Z.Z., Liu Z.Y., Zeng W.B., Cheng S., Ma Y.P., et al. Identification and BAC construction of HAN, the first characterized HCMV clinical strain in China. J Med Virol. 2016; 88 (5): 859–870.

12. Lee Y., Kim M., Han J., Yeom K.H., Lee S., Baek S.H., et al. MicroRNA genes are transcribed by RNA polymerase II. EMBO J. 2004; 23(20): 4051–4060.

13. Fraser K.A., Rice S.A. Herpes simplex virus type 1 infection leads to loss of serine-2 phosphorylation on the carboxy-terminal domain of RNA polymerase II. J Virol. 2005; 79(17): 11323–11334.

14. Fraser K.A., Rice S.A. Herpes simplex virus immediate-early protein ICP22 triggers loss of serine2-phosphorylated RNA polymerase II. J Virol. 2007; 81(10): 5091–5101.

15. Perng Y.C., Campbell J.A., Lenschow D.J., Yu D. Human cytomegalovirus pUL79 is an elongation factor of RNA polymerase II for viral gene transcription. PLoS Pathog. 2014; 28; 10(8): e1004350. doi: 10.1371/journal.ppat.1004350.

16. Hoadley K.A., Yau C., Wolf D.M., Cherniack A.D., Tamborero D., Ng S., et al. Multiplatform analysis of 12 cancer types reveals molecular classification within and across tissues of origin. Cell. 2014; 158:929–944.

17. Franco H.L., Kraus W.L. No driver behind the wheel? Targeting transcription in cancer. Cell. 2015; 163:28–30.

18. Santamaria N.G., Robles C.M., Giraudon C., Martinez-Leal J.F., Compe E., Coin F., et al. Lurbinectedin specifically triggers the degradation of phosphorylated RNA polymerase II and the formation of DNA breaks in cancer cells. Mol Cancer Ther. 2016; 15 (10): 2399–2412.

19. Berbamini G., Reschke M., Battista M.C., Boccuni M.C., Campanini F., Ripalti A., et al. The major open reading frame of the beta2.7 transcript of human cytomegalovirus: in vitro expression of a protein posttrancriptionally regulated by the 5’ region. J Virol. 1998; 72 (10):8425–8429.

20. Jault F.M., Jault J.M., Ruchti F., Fortunato E.A., Clark C., Corbeil J., et al. Cytomegalovirus infection induces high levels of cyclins, phosphorylated Rb, and p53, leading to cell cycle arrest. J Virol. 1995; 69 (11): 6697–6704.

21. Noris E., Zannetti C., Demutas A., Sinclair J., De Andrea M., Gariglio M., et al. Cell cycle arrest by human cytomegalovirus 86-kDa IE2 protein resembles premature senescence. J Virol. 2002; 76(23): 12135–12148.

22. Wohlschlegel J.A., Dwyer B.T., Dhar S.K., Cvetic C., Walter J.C. Inhibition of eukaryotic DNA replication by geminin binding to Cdt1. Science. 2000; 290(5500): 2309–2312.

23. Nishitani H., Taraviras S., Lygerou Z., Nishimoto T. The human licensing factor for DNA replication Cdt1 accumulates in G1 and is destabilizer after initiation of S-phase. J Biol Chem. 2001; 276(48): 44905–44911.

24. Speck C., Chen Z., Li H., Stillman B. ATPase-dependent cooperative bindings of ORC and cdc6 to origin DNA. Nat Struct Mol Biol. 2005; 12(11): 965–971.

25. Randell J.C., Bowers J.L., Rodriguez H.K., Bell S.P. Sequential ATP hydrolysis by cdc6 and ORC directs loading of the Mcm2-7 helicase. Mol Cell. 2006; 21(1): 29–39.

26. Fortunato E.A., Sanchez V., Yen J.Y., Spector D.H. Infection of cells with human cytomegalovirus during S phase results in a blockade to immediate-early gene expression that can be overcome by inhibition of the proteasome. J Virol. 2002; 76: 5369–5379.

27. Braun T.E., Poole E., Sinclair J. Depletion of cellular pre-replication complex factors results in increasing human cytomegalovirus DNA replication. PLos One. 2012; 7(5):e36057. doi. 10.1371/journal.pone.0036957.

28. McSharry B.P., Tomasec P., Neale M.L., Wilkinson G.W. The most abundantly transcribed human cytomegalovirus gene (beta2.7) is non-essential for growth in vitro. J Gen Virol. 2003; 84(Pt9): 2511–2516.

